# Dental abscesses on the maxilla of a two million-year-old early *Homo* specimen

**DOI:** 10.1101/595595

**Authors:** Ian Towle, Joel D. Irish

**Author notes:** **Author contact details**: Joel D. Irish.

## Abstract

Abscesses and other periapical lesions are found in abundance in recent archeological samples, yet are scarce in the fossil hominin record. Periapical voids commonly develop after exposure of a tooth’s pulp chamber and are commonly associated with heavy crown wear, trauma or caries. In this study, all available maxilla and mandible fragments from the South African fossil hominin collections were studied, including specimens assigned to *Homo naledi, Paranthropus robustus, Australopithicus africanus, A. sediba* and early *Homo*. Only one specimen displayed voids consistent with periapical lesions, and a differential diagnosis of these voids was undertaken. The specimen, SK 847, is described as early *Homo* and has been dated to 2.3-1.65MA. There is one definite abscess, and likely at least two more with postmortem damage, all on the anterior aspect of the maxilla and associated with the incisors. The abscesses originate from the apices of the incisor roots and are therefore unlikely to represent a systemic disease such as multiple myeloma. They best fit the description of an abscess rather than a cyst or granuloma, with one showing a rounded thickened rim around the lesion. The abscesses highlight that this individual used their anterior dentition extensively, to the point that the pulp chambers were exposed on multiple teeth. This is one of the earliest hominin examples of a dental abscess and shows that this individual was able to cope with several concurrent abscesses, clearly surviving for an extended period. Therefore, this finding adds additional information to the history of dental pathology in our genus.

## Introduction

Abscesses and other periapical voids are commonly found in the mandible and maxilla of humans and other animals (e.g., Miles & Grigson, 2003; Sauther et al., 2002; Eshed et al., 2006). These voids can develop after exposure of a tooth’s pulp chamber, often through heavy wear (i.e., attrition or abrasion), trauma and caries (Linn et al., 1987; Nair, 2004). As well as abscesses, other types of periapical voids include cysts, fenestrations and granulomata, and concerns have been raised about the incorrect use of these different terms in archaeological studies (Nair, 2004; Dias & Tayles, 1997). In particular, the majority of voids may be painless granulomata, with evidence suggesting that less than a third may be chronic abscesses (Ogden, 2008). However, because a granuloma can become a cyst, both can become an abscess, and many disagree as to what is considered an abscess, there will be times when a lesion is difficult to decipher in the archaeological record.

The frequency of abscessing varies substantially among human populations, as well as non-hominin primate species (Lieverse et al., 2007; Eshed et al., 2006; Cuozzo & Sauther, 2012; Dias & Tayles, 1997; Miles & Grigson, 2003; Schultz, 1956). Abscesses, and other periapical lesions, are commonly associated with both heavy crown wear and carious lesions in recent human archaeological samples. Until recently, they had rarely been reported in the hominin fossil record (Leek, 1966; Lacy et al., 2012; Lacy, 2014; Margvelashvili et al., 2016; Martinón-Torres et al., 2011; Gracia-Téllez et al., 2013). The most important factor in extant primates, as a whole, is severe occlusal attrition (Legge, 2012). Great apes appear to have higher rates than do other primates, with the canine most frequently involved (Schultz, 1956).

In this study, we recorded the presence/absence of periapical voids in South African fossil hominins, for comparison with those reported in more recent populations and other primate species. Our working hypothesis was that their presence likely resulted from crown wear, given its ubiquity in many hominin specimens, rather than caries, which are comparatively uncommon (Towle, 2019; Grine et al., 1990). We then conducted a differential diagnosis on one early specimen of *Homo* presenting several large voids on the anterior aspect of the maxilla to discern the etiology.

## Materials and methods

All maxilla and mandible fragments available for study in the South African fossil hominin collections were observed, including specimens assigned to *Homo naledi, Paranthropus robustus, Australopithicus africanus, A. sediba* and Early *Homo*. For a full specimen list and species classifications see Towle (2017). Specimens are curated at The Ditsong National Museum of Natural History and University of the Witwatersrand. Lesions were photographed using a Dino-Lite^®^ camera (Dino-Lite AM2111 handheld microscope, 10x to 50x zoom magnification).

Any antemortem bone voids present in a maxilla or mandible associated with a tooth root apex were recorded. If a lesion was not associated with an apex it is not included here, since a systemic disease such as multiple myeloma and metastatic carcinoma can be responsible (Ogden, 2008; Odes et al., 2018). A tooth had to have all the surrounding bone present to be included. Only one specimen met these criteria to conduct a differential diagnosis. Methods suggested by Dias and Tayles (1997) and Ogden (2008) were used, firstly to describe the lesions and secondly to suggest an etiology.

## Results and discussion

The specimen, SK 847, is described as early *Homo* dated to 2.3-1.65MA (Tobias, 1991). Although the phylogenic placement and associated material of this specimen has been debated, it is commonly placed in the genus *Homo* (e.g., Berger et al., 2015; Grine et al., 1993). It is usually referred to as SK 847, after two fragments were incorporated (Clarke & Howell, 1972; Grine et al., 1993). As well as the maxilla, the specimen also has much of the frontal cranium present (Figure 9.1a) and has been associated with different mandibles (Clarke & Howell, 1972; Grine et al., 1993). Only the left lateral maxillary incisor remains, with severe wear evident (wear grade 8 in Smith, 1984; Figure 9.1c). In-depth descriptions of the specimen are available in Robinson (1953) and Clarke (1979).

This individual likely had at least three abscesses at the time of death—all associated with the incisors (Figure 9.1d). Two evidence post-mortem damage; however, this likely relates to an already existing void, given the definite abscess associated with the left upper central incisor (Figure 9.1e). The left upper central incisor is absent, likely post-mortem, judging by the lack of alveolar resorption. There is also a potential abscess on the lingual surface, from what could have been a drainage channel associated with right lateral incisor. However, this void may instead be an unusual foramen, as noted by Robinson (1953). The heavy wear on the remaining teeth, in particular the removal of most enamel on the left lateral incisor, suggests the abscesses resulted from pulp chamber perforation. The abscesses originate from the apices of the incisor roots, so are unlikely to result from a systemic disease such as multiple myeloma (Dias & Tayles, 1997). The void best fits the description of an abscess rather than cyst or granuloma, with a rounded thickened rim around the void (Figure 1.1e; Dias & Tayles, 1997; Ogden, 2008).

**Figure 1.**
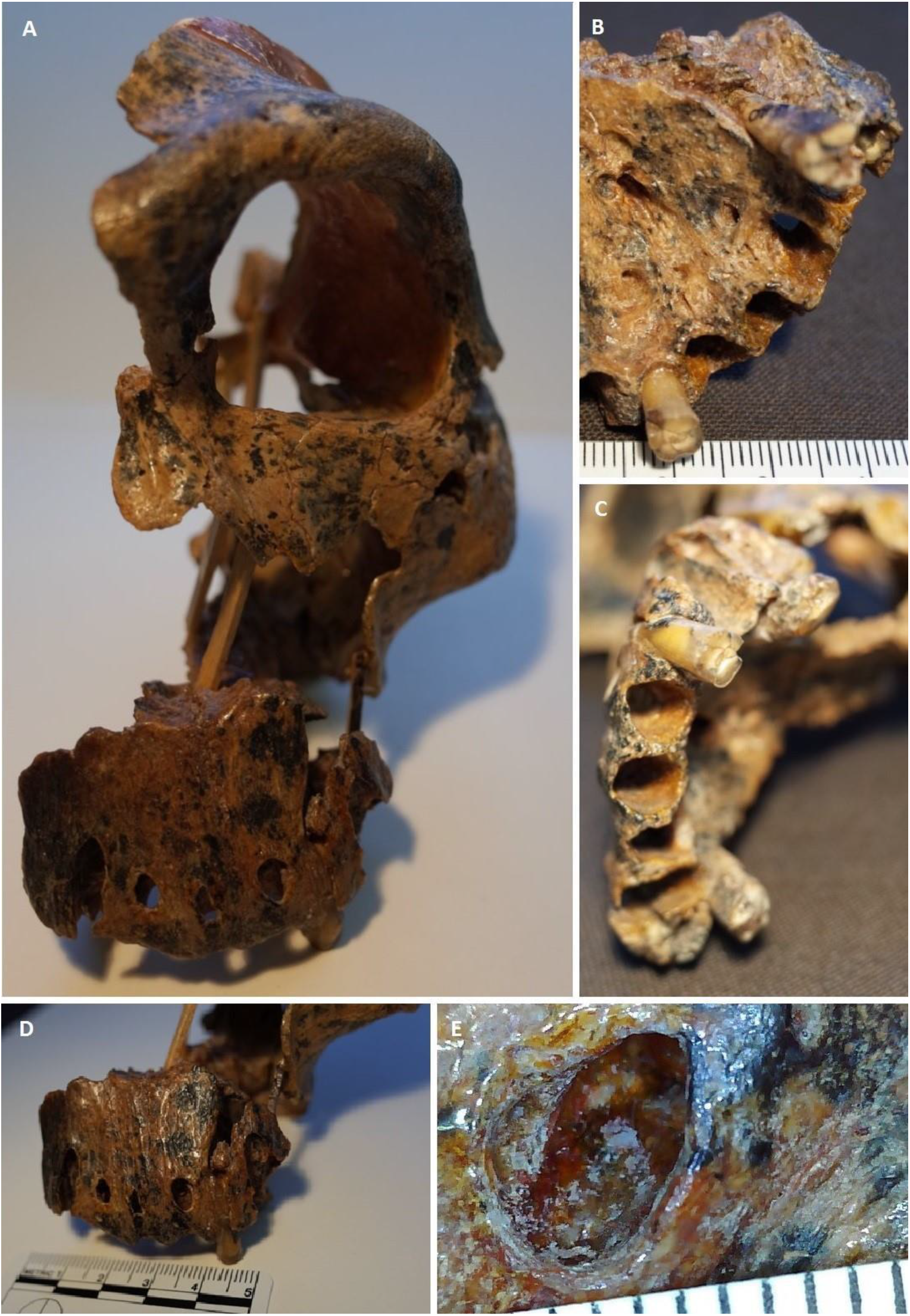
Specimen SK 847 (early *Homo):* A) overview of the entire specimen; B) lingual aspect; C) heavy wear on left lateral upper incisor; D) buccal abscesses; E) detail of buccal abscess, showing thickened bone around the circumference of the abscess.

Robinson (1953) first described this specimen, which at the time consisted only of the maxilla (SK 80), with a focus on its morphology; the extreme incisor wear was also noted and described. Clarke (1979) concentrated on the morphology as well, though described the state of preservation and maxillary voids. He did not mention whether the latter were ante- or postmortem in nature, but reported periodontitis on the lingual side of the right lateral incisor (Clarke, 1979). Thus, the present study builds on this research and suggests the individual suffered from several, connected dental issues, i.e., extreme wear, periodontal disease and dental abscesses.

Extreme wear removed the enamel on the anterior teeth and much of the dentine, which ultimately led to exposure of the root canals. Subsequently, based on clinical progression, periapical periodontitis would have occurred, caused by chronic root canal infection. Eventually, several periapical cysts would have formed in the anterior maxilla of this individual’s mouth and ultimately to abscesses (Dias & Tayles, 1997). This process is common in modern humans, with maxillary anterior teeth most affected (Nair, 2004). Similar lesions caused by heavy wear have been found in archaeological samples of hunter-gatherers (Lieverse et al., 2007; Eshed et al., 2006), as well as in extant primates (Cuozzo & Sauther, 2012). Abscesses and other periapical voids have rarely been reported in fossil remains, but recent studies suggest they may not be as uncommon as once thought (e.g., Lacy et al., 2012; Lacy, 2014; Margvelashvili et al., 2016; Martinón-Torres et al., 2011; Gracia-Téllez et al., 2013). In particular, such voids—again likely related to wear—were described in another specimen of early *Homo* (i.e., Mandible D2600 from Dmanisi), which potentially belongs to the same species, *Homo erectus*, and is roughly contemporary with SK 847, dating to 1.77MA (Margvelashvili et al., 2016). Periapical voids have been more frequently described in later *Homo* (Gracia-Téllez et al., 2013; Rosas et al., 2006; Lebel et al., 2001), though not all were wear related, e.g., Kabwe 1 (Lacy, 2014).

In sum, the voids in the SK 847 maxilla are of interest on several levels. First, at 2.3-1.65MA, this may be the earliest example of dental abscessing in fossil hominins yet known. Second, the number and severity of these abscesses suggest that the individual suffered with them for some time. That said, third, their severity and unhealed state might also indicate they contributed to the individual’s death, i.e., septicemia (Gracia-Téllez et al., 2013). Indeed, given the increased prevalence of abscesses through time, others in the genus *Homo* may have been affected similarly. Finally, the lack of any abscesses in the remaining South African fossil hominin specimens inspected for this study, including large samples of *P. robustus* and *A. africanus*, suggest this individual, and perhaps early *Homo* on a broader scale, practiced a novel behavior. S/he may have regularly processed abrasive food items with her/his incisors (i.e., attrition) or, perhaps, used them for a non-masticatory activity, such as a ‘third hand’ (in which case the wear would be characterized as abrasion). In any event, this and alternate evidence of dentition-related behavior in the South African collections, e.g., anterior root grooves in an *A. africanus* specimen (Towle et al., 2018), and high rates of enamel fractures in *H. naledi* (Towle et al., 2017), help personalize the study of our hominin ancestors and relatives. In this case, the SK 847 abscesses provide some measure of individuality to this once-living individual.

## Acknowledgements

The authors thank L. Berger and B. Zipfel from the University of the Witwatersrand and S. Potze from the Ditsong Museum of South Africa for access to their collections. This research was supported by a studentship to the first author from Liverpool John Moores University.

